# Characteristics of viable but nonculturable *Vibrio parahaemolyticus* induced under extended periods of cold-starvation with various NaCl concentrations

**DOI:** 10.1101/2020.07.14.202127

**Authors:** Jae-Hyun Yoon, Jeong-Eun Hyun, Sung-Kwon Moon, Sun-Young Lee

## Abstract

This study was undertaken to examine the induction of VBNC states of *Vibrio parahaemolyticus* under prolonged cold-starvation with various NaCl concentrations and their responsive characteristics to maintain cell viability. *V. parahaemolyticus* entered the viable but nonculturable (VBNC) state in artificial sea water at 4°C within 80 day and persisted in the VBNC state for 150 days. During cold-starvation, bacterial cells were used to estimate their cell functions, including cytotoxicity, fatty acid composition, membrane potential, and morphology. Cytotoxic effect of *V. parahaemolyticus* cells against animal cell lines was decreased to below 50% after 80 days. VBNC *V. parahaemolyticus* cells showed decreasing levels of palmitic, vaccenic, and hexadecenoic acid on membrane, concomitantly with the formation of empty gaps between the cytoplasmic and outer membrane, in comparison with those of the pure cultures. Starvation at 4°C for 30 days resulted in a high increase in _N_-phenyl-_1_-napthylamine intensity within *V. parahaemolyticus* cells. Membrane potential and cellular composition were strongly affected by increasing NaCl contents of the microcosms after its evolution into the VBNC state. VBNC *V. parahaemolyticus* cells may undergo selected physiological changes such as the modulation of membrane potential and re-arrangement of cellular composition.

## Introduction

*Vibrio parahaemolyticus* has been found in marine environments and readily isolated from a wide variety of raw aquatic products such as clam, mussel, oyster, scallop, and shrimp during warmer season, when the incidence of food-borne disease outbreaks is the highest (1–4). Consumption of marine products contaminated with *V. parahaemolyticus* results in various clinical symptoms, ranging from acute abdominal pain, vomiting, and nausea to fatal septicemia (5). Especially, major food-borne pathogens, such as *Campylobacter jejuni*, *Escherichia coli* O157:H7, *Shigella dysenteriae V. parahaemolyticus*, and *Vibrio vulnificus*, are known to become viable but nonculturable (VBNC) under various environmental stresses, including CO_2_, copper, low temperature, and starvation (6–8). In this physiological state, VBNC microorganisms cannot be cultured on routine media which normally support their growth and may represent specific changes in their cell morphology (9), fatty acid (10) and protein profiles (11), and metabolic activity (12), thereby maintaining Miles their cell membrane integrity in response to various hostile environments. VBNC bacteria are also characterized by reduced cell metabolic activities, including ATP synthesis, cytoplasmic volume, diffusion of macromolecular components, gene expression, production of metabolite, and transcript (13, 14).

Especially, low temperature (5-10°C) in combination with nutrient-deprivation was shown to be strongly involved in the loss of culturability and entry of *V. parahaemolyticus* into the VBNC state. Bates and Oliver (15) showed that when Kanagawa positive strains of *V. parahaemolyticus* were incubated in artificial sea water (ASW) at 5°C for ≤20 days, these organisms were undetectable but viable at levels of 10^3-6^ CFU/ml, as determined by epifluorescence microscopy with acridine orange. In a study conducted by Wong et al. (1), *V. parahaemolyticus* ST550 was induced into the VBNC state in Morita mineral salt solution (MMS) at 4°C for 32 days. Previous studies indicate that approximately 30-70 days were required for *V. parahaemolyticus* strains to enter the VBNC state at 3-5°C (10, 16, 17). Importantly, conversion of VBNC forms can be accelerated more rapidly when *V. parahaemolyticus* cells in the stationary growth phase were incubated in ASW microcosms supplemented with high concentrations of NaCl at 4°C (18, 19). Cold-starvation in ASW microcosms supplemented with 5%, 10%, and 30% NaCl at 4°C led to the phase transition of *V. parahaemolyticus* into the VBNC state within 30, 14-21, and 3 days, respectively. Considering that food preservation processing such as low pH or high NaCl are commonly used to prevent the growth of spoilage and pathogenic organisms on food, the addition of NaCl may shorten the incubation-times that are needed for *V. parahaemolyticus* to enter the VBNC state. As microorganisms in such a dormant but viable state may be recovered in a favorable environment where provides sufficient energy sources to encourage their biological function and growth, the incidence of VBNC pathogens on food would be closely involved in the food-borne disease outbreaks and pose a potential risk to public health. However, little is known about how high NaCl contents in nutrient-deficient microcosms affect the formation and physiological characteristics of VBNC *V. parahaemolyticus* induced by prolonged cold-starvation. Exploring cell properties of *V. parahaemolyticus* in response to various environmental conditions may be critical for better understanding the ecology of this pathogen, as well as its survival mechanisms. In this study, *V. parahaemolyticus* ATCC 17802, *V. parahaemolyticus* ATCC 33844, and *V. parahaemolyticus* ATCC 27969 were incubated in ASW microcosms (pH 6) supplemented with various NaCl concentrations at 4°C until these bacteria were induced into the VBNC state. Physiological properties of VBNC *V. parahaemolyticus* were characterized by measuring cytotoxic effects to animal cell lines, membrane potential with _N_-phenyl-_1_-napthylamine (NPN) uptake, intracellular leakage of nucleic acid and protein, cell hydrophobicity, and morphological change. Additionally, fatty acid composition on cell membrane of *V. parahaemolyticus* was analyzed before and after the induction of VBNC cells.

## Materials and methods

### Preparation of microcosm

According to the instructions provided by a reliable supplier, 30 g of sea salt powder (Sigma-Aldrich, St. Louis, MO, USA) consisting of 19,290 mg of Cl, 10,780 mg of Na, 2,660 mg of SO_4_, 420 mg of K, 400 mg of Ca, 200 mg of CO_3_, 8.8 mg of Sr, 5.6 mg of B, 56 mg of Br, 0.24 mg of I, 0.3 mg of Li, 1.0 mg of F, and 1,320 mg of Mg, were dissolved in 1 l of distilled water (DW). To determine effects of higher NaCl concentrations on the induction of *V. parahaemolyticus* into the VBNC state, each of microcosms was modified by adding excessive amounts of NaCl (5%, 10%, and 30%) and its acidity was adjusted to pH 6 using membrane-filtered 1 N NaOH (Kanto chemical, Tokyo, Japan). The modified microcosms had different NaCl contents: 0.75% (ASW), 5% (ASW_5_), 10% (ASW_10_), and 30% NaCl (ASW_30_). All microcosms were autoclaved at 121°C for 20 min prior to use.

### Preparation of bacterial inoculums

*V. parahaemolyticus* ATCC 17802, *V. parahaemolyticus* ATCC 33844, and *V. parahaemolyticus* ATCC 27969 were purchased from the Korean Collection for Type Cultures (KCTC, Daejon, Korea). Each bacterial stock was maintained at −75°C and was activated in tryptic soy broth (Difco, Detroit, MI, USA) added with 3% NaCl (TSB_S_) at 37°C for 24 h. *V. parahaemolyticus* cells in the stationary phase were harvested by centrifugation at 10,000 × g for 3 min, washed in ASW, and the final pellets were re-suspended in 1 ml of ASW, corresponding to approximately 10^8-9^ CFU/ml. Each bacterial suspension was inoculated into ASW, ASW_5_, ASW_10_, and ASW_30_, respectively. The microcosms were kept at 4°C until the culturable counts of *V. parahaemolyticus* decreased to below the detection limits (< 1.0 log CFU/ml).

### Enumeration

*V. parahaemolyticus* ATCC 17802, *V. parahaemolyticus* ATCC 33844, and *V. parahaemolyticus* ATCC 27969 were plate-counted on tryptic soy agar (Difco) amended with 3% NaCl (TSA_S_). Decimal dilutions (10^−1^) were prepared in alkaline peptone water (APW, Difco) consisting 10 g of peptone and 10 g of NaCl in 1 l of sterile DW. Then, 100 μl of these aliquots were spread on TSA_S_. Each agar plate was incubated at 37°C for 24 h, and colonies shown on the media were enumerated.

### Epifluorescence microscopy with SYTO_9_ and propidium iodide

Total and viable counts of *V. parahaemolyticus* were measured using the Live/Dead® BacLight™ Bacterial Viability Kit (Invitrogen, Mount Waverley, Victoria, Australia) containing two nucleic acid stains, SYTO9 and propidium iodide (PI). While SYTO9 has a high affinity for DNA and chromosome and is used for labelling bacterial cells with intact and compromised membranes, PI selectively penetrates bacterial cells with damaged membranes. Briefly, equal volumes (1:1) of SYTO9 and PI were combined, and 3 μl of this mixture were added to 1 ml of the bacterial suspension. After 15 min of incubation at 25°C in the dark, 5-8 μl of the bacterial aliquots were attached on a glass slide. Bacterial images were demonstrated via an electron-fluorescent microscope (TE 2000-U, Nikon, Tokyo, Japan).

### Cytotoxicity assay

Caco2 and Vero cell lines were cultured in 5-10 ml of Dulbecco’s modified eagle’s medium (DMEM, Corning, NY, USA) supplemented with 5% (DMEM_5_) and 20% (DMEM_20_) fetal bovine serum (FBS, Corning) at 37°C for 2 days in 5% CO_2_, respectively. After 2 days of incubation, each DMEM solution was removed in a petri-dish and washed in 5 ml of PBS three times. Each culture was added by 5 ml of trypsin (Corning) for cell lysis and incubated at 37°C for 5 min in 5% CO_2_. To alleviate the enzymatic activity caused by trypsin, 2-3 ml of the DMEM media, such as DMEM_5_ and DMEM_20_, were added to Caco2 and Vero cells, respectively. Animal cell fluids were further transferred to sterile cap tubes and centrifugated at 15,000 × g for 3 min. The supernatants were eliminated, and cell pellets from Caco2 and Vero were re-suspended, corresponding to the cell density of 10^4^ ml^−1^, in 5 ml of DMEM_5_ and DMEM_20_, respectively. Then, 100 μl of Caco2 and Vero were loaded into 96-well plates containing 100 μl of DMEM_5_ and DMEM_20_, respectively. The eukaryotic cell lines were incubated at 37°C for 24 h in 5% CO_2_ before use. At regular time-intervals, *V. parahaemolyticus* cells incubated in ASW microcosms were withdrawn from the incubator. The bacterial aliquots (100 μl) were added to 96-well plates containing 100 μl of each cell lines and were incubated at 37°C for 24 h in 5% CO_2_. Five mg ml^−1^ of 3-(4, 5-dimethylthiazol-2-yl)-2, 5 diphenyl tetrazolium bromide (MTT, Corning) was added to each well in the 96-well plates, and the cell fluids were incubated at 37°C for 1 h. The culture fluids were added by 100 μl of DMSO (Corning) and were read on a microtiter plate reader at optical densities (ODs) between 570 and 620 nm (Multiskan GO Microplate Spectrophotometer, Thermo Scientific, Vantaa, Finland).

### Measurement of membrane potential

Biological function of cell membrane was determined by NPN uptake assay. At regular time-intervals, *V. parahaemolyticus* cells incubated in ASW microcosms were withdrawn from the incubator. The bacterial solutions (1 ml) were centrifugated at 15,000 × g for 3 min, washed in 4-(2-hydroxyethyl)-1-piperazineethanesulfonic acid (HEPES; Thermo Fisher Scientific, Waltham, MA, USA) two times, and were re-resuspended in 2 ml of HEPES. NPN was added to the mixtures, corresponding to the final concentration of 10 μM, and the background fluorescence was recorded via a spectrophotometer (Gemini XPS, Molecular Devices Inc., CA, USA). Excitation and emission wavelengths were adjusted at 350-420 nm. Polymyxin B (Sigma) was used as a positive control due to its outer membrane permeabilizing property.

### Leakage of cellular components

Bacterial solutions (1.5 ml) of *V. parahaemolyticus* incubated in ASW, ASW_5_, ASW_10_, and ASW_30_ at 4°C for 100 days were transferred to sterile microtubes and were centrifugated at 15,000 × g for 3 min. Each cell supernatant was collected and used to assess the leakage of cellular components such as DNA and protein via a microtiter plate reader (Multiskan GO Microplate Spectrophotometer, Thermo Scientific) at OD_570 nm_ and OD_620 nm_, respectively.

### Measurement of enzymatic activity

Catalase activity was measured, using a spectrophotometric H_2_O_2_-degradation assay (CAT100, Sigma-Aldrich). Briefly, the pure cultures of *V. parahaemolyticus* or VBNC *V. parahaemolyticus* cells incubated in ASW microcosms at 4°C for 100 days were re-suspended in 50 mM potassium phosphate buffer (pH 7) containing 1 g of 3-mm-sized glass bead (Sigma), vortexed for 25 min, and were centrifugated at 15,000 × g for 3 min. The supernatants were separately transferred to sterile microtubes. In a total of 100 μl of volume, 15 μl of the supernatant were mixed with 5 mM H_2_O_2_ and incubated at 25°C for 15 min. This reaction ceased by the addition of 900 μl of 15 mM sodium azide. The absorbance was colorimetrically read at 520 nm via a multi-scan Go spectrophotometer (Thermo Scientific Inc.).

### Cell hydrophobicity

One ml of *V. parahaemolyticus* grown overnight in TSB_S_ at 37°C and incubated in ASW microcosms at 4°C for 50 days were centrifugated at 15,000 × g for 3 min, washed in PBS twice, and were re-suspended in PBS to fit an OD of 1.0 (A_o_) at 600 nm via a UV-Visible Spectrometer (Multiskan GO Microplate Spectrophotometer, Thermo Scientific). One hundred μl of hexadecane was added to 1 ml of the bacterial solution and was incubated at an ambient temperature for 10 min. ODs of the mixtures in aqueous phase were measured at 600 nm (A_1_). The degree of hydrophobicity was calculated, following as[1-A_1_/A_o_] ×100 (%).

### Transmission electron microscopy (TEM)

*V. parahaemolyticus* cells grown in TSB_S_ at 37°C for 24 h and incubated in ASW and ASW_5_ at 4°C for 100 days were centrifugated at 15,000 × g for 3 min, rinsed in 0.1M PBS (pH 7) three times, and were re-suspended in 0.1M PBS, respectively. The cell fluids were pre-fixed in 2% paraformaldehyde overnight at 4°C. The cell solutions were washed in 0.1M PBS, post-fixed in 1% osmium tetroxide, and were serially dehydrated by 30%, 50%, 70%, 95%, and 100% ethanol solutions. Each of them was infiltrated with 2 ml of epoxy resin. Polymerization of the resins was performed at 60°C for 24 h. The resins were cut (section: approximately 120 nm thickness) and were photographed with a JEOL JEM 1200 EX transmission electron microscope (JEOL USA Inc., Peabody, MA, USA).

### Fatty acid composition

Fatty acid analysis was carried out according to the standard protocol provided by the Microbial Identification System (MIDI, Microbial ID Inc., Newark, Del., USA). *V. parahaemolyticus* was harvested by centrifugation at 15,000 × g for 3 min and was processed by saponification, methylation, and extraction of carboxylic acid derivatives from long-chain aliphatic molecules. The extracted lipids were analyzed by gas chromatography (GC) and identified, using the TSBA6 database of the MIDI system.

## Results

### Bacterial counts

Fig 1 represents the survival (log CFU/ml) curves of *V. parahaemolyticus* ATCC 17802, *V. parahaemolyticus* ATCC 33844, and *V. parahaemolyticus* ATCC 27969 incubated in ASW (pH 6) microcosms containing various NaCl concentrations at 4°C for 90 days. The initial cell densities of *V. parahaemolyticus* ranged from 6.0 to 8.0 log CFU/ml. *V. parahaemolyticus* ATCC 17802 exceeded approximately 4.0 log CFU/ml when incubated in ASW and ASW_5_ at 4°C for 40 days, whereas these bacteria dropped to below the detection limits (< 1.0 log CFU/ml) after 80 days. *V. parahaemolyticus* ATCC 17802 became undetectable in ASW_10_ and ASW_30_ at 4°C within at least 21 days. Similarly, *V. parahaemolyticus* ATCC 33844 and *V. parahaemolyticus* ATCC 27969 declined slowly in ASW and ASW_5_ during the first 28 days at 4°C, but these cells remained culturable until day 42. *V. parahaemolyticus* strains were also uncultivable in ASW_10_ and ASW_30_ due to the lack of its culturability on TSA_S_ on day 21. Importantly, the times needed for the complete loss of culturability were lessened with the increasing amounts of NaCl in cold-starvation conditions. While all *V. parahaemolyticus* strains took ≥2 months in ASW to become the VBNC state, approximately 3-21 days were required to do so in ASW_30_. *V. parahaemolyticus* ATCC 17802 dropped to the detection limits in ASW, ASW_5_, ASW_10_, and ASW_30_ at 4°C for 60, 60, 21, and 4 days, respectively.

**Fig 1.** Loss of the culturability and viability of *Vibrio parahaemolyticus.* Culturability (A, C, and E) and viability (B, D, and F) of *Vibrio parahaemolyticus* ATCC 17802 (A-B), *V. parahaemolyticus* ATCC 33844 (C-D), and *V. parahaemolyticus* ATCC 27969 (E-F) in artificial sea water (ASW; pH 6) microcosms at 4°C. •, ASW; ○, ASW_5_; ▾, ASW_10_; ▵, ASW_30_; black bar, initial cell number at 4°C for 0 day; grey bar, total cell number at 4°C for 100 days; intact cell number at 4°C for 100 days.

### Cytotoxicity

Fig 2 shows the cytotoxic effects of VBNC *V. parahaemolyticus* on the survivals of Caco2 and Vero cells. Initially, the animal cell lines were completely disrupted, following co-culture with three strains of *V. parahaemolyticus* grown overnight in TSB_S_ at 37°C. *V. parahaemolyticus* ATCC 17802 remained highly virulent in ASW consistently for killing more than 95% of Caco2 and Vero cell lines (Fig 2). *V. parahaemolyticus* ATCC 33844 exhibited the decreasing cytotoxic effects to Vero cells at levels of 100%, 63%, 82%, and 39% when maintained in ASW at 4°C for 0, 7, 21, and 80 days. There were gradual decreases in the cytotoxic activities of *V. parahaemolyticus* ATCC 27969 on the inactivation of Vero cells with the prolonged incubation periods under cold-starvation conditions (*data not shown*). After resuscitation process, *V. parahaemolyticus* cells were transformed to a culturable state in TSB_S_, concomitantly with the recovered cytotoxic activities between 50-100% more than those of these bacteria exposed to cold-starvation until 80 days. In addition, there were no differences in the cytotoxic effects of VBNC *V. parahaemolyticus* cells with regard to the increasing NaCl contents of the ASW microcosm.

**Fig 2.** Evaluation of cytotoxicity of *Vibrio parahaemolyticus*. *V. parahaemolyticus* ATCC 17802 (A and D), *V. parahaemolyticus* ATCC 33844 (B and E), and *V. parahaemolyticus* ATCC 27969 (C and F) incubated in artificial sea water (ASW; pH 6) microcosms at 4°C for 0, 7, 21, and 80 days against Caco2 (A-C) and Vero (D-F) cells. *, not determined.

### Membrane potential and cellular leakage

Outer membrane permeabilizing activity in VBNC *V. parahaemolyticus* cells was determined using the NPN assay in Table 1. VBNC *V. parahaemolyticus* ATCC 17802 showed the increased fluorescence intensities between 1,769 and 1,875 in ASW_5_, ASW_10,_ and ASW_30_, as compared with those of the actively growing cells. It was shown that 30 days of starvation at 4°C caused the high increases in the fluorescence intensity within VBNC *V. parahaemolyticus* cells. In common, *V. parahaemolyticus* strains yielded the highest NPN uptakes at levels of 1,875-2,643 in ASW_10_ after 30 days. Furthermore, subsequent cellular leakages were observed in *V. parahaemolyticus* cells before and after 150 days of cold-starvation stress (Table 2). Initially, DNA and protein leakages of *V. parahaemolyticus* ATCC 33844 were 0.576-1.562 at 260 nm and 0.466-1.316 at 240 nm, respectively. DNA and protein were found to be released at levels of 2.315-2.683 at 260 nm and 1.218-1.433 at 240 nm from *V. parahaemolyticus* cells exposed to cold-starvation on day 150, respectively. Regardless of the bacterial strains used, the cellular leakages from VBNC *V. parahaemolyticus* cells were increased with the increasing NaCl concentrations of the ASW microcosms.

**Table 1.** Measurement of _N_-phenyl-_1_-napthylamine (NPN) uptake (RFU) of *Vibrio parahaemolyticus* before and after incubation in artificial sea water (ASW, pH 6) microcosms at 4°C for 30 days.

| *V. parahaemolyticus*<br>strain | TSB<sub>S</sub> * | TSB<sub>S</sub>+PB † | Microcosm ‡ |  |  |  |
| --- | --- | --- | --- | --- | --- | --- |
|  |  |  | ASW | ASW<sub>5</sub> | ASW<sub>10</sub> | ASW<sub>30</sub> |
| ATCC 17802 | 1,710 | 1,922 | 1,701 | 1,769 | 1,875 | 1,785 |
| ATCC 33844 | 1,962 | 2,581 | 2,334 | 2,284 | 2,530 | 2,404 |
| ATCC 27969 | 1,737 | 2,810 | 2,295 | 2,533 | 2,643 | 2,068 |

\* Each *V. parahaemolyticus* was grown in tryptic soy broth supplemented with 3% NaCl at 37°C for 24h (control).
† Each *V. parahaemolyticus* was grown in tryptic soy broth added with 3% NaCl and polymyxin B (PB) at 37°C for 24h. (positive control).
‡ Stationary growth phase cells of *V. parahaemolyticus* were incubated in the modified microcosms with different NaCl contents; 0.75% (ASW), 5% (ASW<sub>5</sub>), 10% (ASW<sub>10</sub>), and 30% NaCl (ASW<sub>30</sub>) at 4°C.

**Table 2.** Measurement of intracellular leakages (nucleic acid by OD_260 nm_ and protein by OD_240 nm_) of *Vibrio parahaemolyticus* before and after incubation in artificial sea water (ASW, pH 6) microcosms for 150 days at 4°C.

|  |  | Microcosm * |  |  |  |  |  |  |  |
| --- | --- | --- | --- | --- | --- | --- | --- | --- | --- |
| *V. parahaemolyticus* strain |  | ASW |  | ASW<sub>5</sub> |  | ASW<sub>10</sub> |  | ASW<sub>30</sub> |  |
|  |  | Before storage | After storage | Before storage | After storage | Before storage | After storage | Before storage | After storage |
| DNA leakage<br>(OD<sub>260 nm</sub>) | ATCC 17802 | 0.433 | 0.868 | 0.502 | 1.200 | 0.598 | 0.999 | 0.974 | 1.053 |
|  | ATCC 33844 | 0.576 | 2.611 | 0.685 | 2.550 | 0.783 | 2.315 | 1.562 | 2.683 |
|  | ATCC 27969 | 0.751 | 3.255 | 0.778 | 3.417 | 0.659 | 3.385 | 1.107 | 3.755 |
| Protein leakage<br>(OD<sub>240 nm</sub>) | ATCC 17802 | 0.390 | 0.426 | 0.435 | 0.602 | 0.465 | 0.600 | 0.797 | 0.638 |
|  | ATCC 33844 | 0.466 | 1.270 | 0.537 | 1.218 | 0.630 | 1.226 | 1.316 | 1.433 |
|  | ATCC 27969 | 0.573 | 1.588 | 0.597 | 1.747 | 0.521 | 1.872 | 0.910 | 2.351 |

\* Stationary growth phase cells of *V. parahaemolyticus* were incubated in the modified microcosms with different NaCl contents; 0.75% (ASW), 5% (ASW<sub>5</sub>), 10% (ASW<sub>10</sub>), and 30% NaCl (ASW<sub>30</sub>) at 4°C.

### Enzymatic activity

After 100 days of cold-starvation, the catalase activity (U/mg) of *V. parahaemolyticus* was measured as shown in Table 3. The catalase activity of *V. parahaemolyticus* ATCC 17802 was 5.869 U/mg when grown in TSB_S_ at 37°C for 24 h. In contrast, VBNC *V. parahaemolyticus* ATCC 17802 were estimated to approximately 29.343, 51.350, 16.138, and 16.138 U/mg in ASW, ASW_5_, ASW_10_, and ASW_30_, respectively. However, VBNC *V. parahaemolyticus* ATCC 27969 displayed the lower catalase activities at levels of 10.27-24.941 U/mg in ASW, ASW_5_, and ASW_10_ in comparison with the actively growing cells (48.415 U/mg). Nevertheless, *V. parahaemolyticus* ATCC 27969 was able to achieve the highest catalase activity of 111.502 U/mg in ASW_30_ after 100 days. The ability of *V. parahaemolyticus* to hydrolyze reactive oxygen species (ROS) compounds may be dependent on the bacterial strains used and the length of cold-starvation stress, rather than the different NaCl concentrations.

**Table 3.** Catalase activity (U mg^−1^) of *Vibrio parahaemolyticus* in the viable but nonculturable state induced in artificial sea water (ASW, pH 6) microcosms for 100 days at 4°C.

| *V. parahaemolyticus* |  | Microcosm † |  |  |  |
| --- | --- | --- | --- | --- | --- |
| strain | TSB<sub>s</sub> * | ASW | ASW<sub>5</sub> | ASW<sub>10</sub> | ASW<sub>30</sub> |
| ATCC 17802 | 5.869 | 29.343 | 51.350 | 16.138 | 16.138 |
| ATCC 33844 | 14.671 | 5.869 | 45.481 | 19.073 | 63.087 |
| ATCC 27969 | 48.415 | 10.270 | 23.474 | 24.941 | 111.502 |

\* Each *V. parahaemolyticus* was grown in tryptic sob broth supplemented with 3% NaCl at 37°C for 24h (control).
† Stationary growth phase cells of *V. parahaemolyticus* were incubated in the modified microcosms with different NaCl contents; 0.75% (ASW), 5% (ASW<sub>5</sub>), 10% (ASW<sub>10</sub>), and 30% NaCl (ASW<sub>30</sub>) at 4°C.

### Fatty acid composition

Table 4 shows the changes of fatty acid profile in VBNC *V. parahaemolyticus* ATCC 17802 persisted at 4°C for 90 days. Palmitic acid (_C16_, 25.6%) and palmitoleic acid (_C16:1 w7c_, 26.8%) were the most abundant in *V. parahaemolyticus* ATCC 17802 cells grown overnight in TSB_S_. When *V. parahaemolyticus* ATCC 17802 were incubated in ASW microcosms at 4°C for 90 days, palmitic acid was increased slightly, ranging from 23.1% to 23.9%. Among saturated fatty acids, the contents of lauric acid (_C12_), _2_-hydroxylauric acid (_C12 2OH_), and myristic acid (_C14_) were increased as *V. parahaemolyticus* ATCC 17802 became VBNC. Additionally, VBNC cells exhibited the increasing concentrations of palmitoleic acid, ranging from 33.0% to 35.8%. After 90 days, the levels of cis-vaccenic acid (_C18:1 w7c_) were 18.1%, 14.9%, 14.2%, and 13.2% in ASW, ASW_5_, ASW_10_, and ASW_30_, respectively. VBNC *V. parahaemolyticus* ATCC 17802 were restored to a culturable state, following enrichment in a nutrient-rich medium (TSB_S_) at 25°C for 7 days (*data not shown*). In the recovered cells, the total density of unsaturated fatty acid was increased more than that for the VBNC cells. While lauric acid, _2_-ydroxylauric acid, and unknown (_C14 3OH_) were increased, palmitic acid was decreased remarkably among the total concentrations of saturated fatty acid in the recovered cells. In addition, cis-vaccenic acid was also increased largely, ranging from 20.65% to 32.17% after a stress relief. Interestingly, it was found that some fatty acids, such as _3_-hydroxy-_9_-methyldecanoic acid (C_11 iso 3OH_), cetyl alcohol (_C16 N alcohol_), and _cis_-_11_-palmitoleic acid (C_16:1_ w5c), were synthesized exclusively with the increasing NaCl concentrations during cold-starvation. Cetyl alcohol and _cis_-_11_-palmitoleic acid were newly formed when *V. parahaemolyticus* ATCC 17082 persisted in ASW microcosms containing ≥5% NaCl at 4°C for 90 days. On the other hand, palmitic acid, _(7Z)_-_13_-methyl-_7_-hexadecenoic acid (C_17:1 anteiso_), and _cis_-Vaccenic acid were increased by the decreasing NaCl amounts of the ASW microcosms. The results indicate that high NaCl concentrations may be an important factor for inducing an alteration in fatty acid profile of *V. parahaemolyticus* exposed to cold-starvation conditions.

**Table 4.**
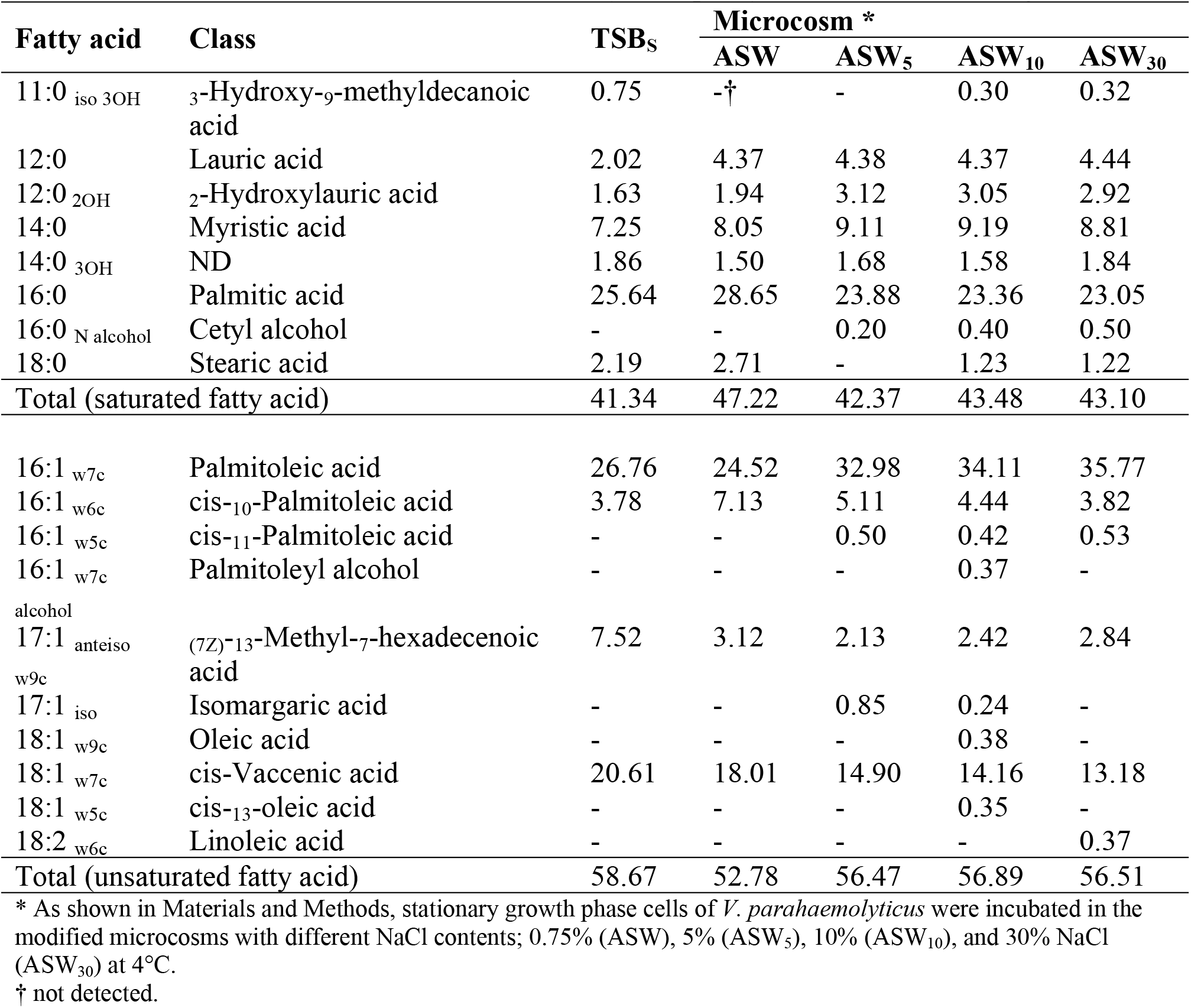
Fatty acid composition (%) of *Vibrio parahaemolyticus* ATCC 17802 the in viable but nonculturable state induced in artificial sea water (ASW, pH 6) microcosms for 90 days at 4°C.

### Cell hydrophobicity

Table 5 depicts the levels of hydrophobicity in VBNC V*. parahaemolyticus* induced at 4°C for 100 days. The initial levels of hydrophobicity were 22.0-37.1, 22.8-26.3, 22.2-45.8, and 20.8-29.8 in ASW, ASW_5_, ASW_10_, and ASW_30_, respectively. VBNC *V. parahaemolyticus* showed the remarkably increasing levels of hydrophobicity more than those for the pure cultures. In ASW_5_, ASW_10_, and ASW_30_, the hydrophobic properties of *V. parahaemolyticus* ATCC 17802 exposed to cold-starvation for 100 days were 37.5%, 46.8%, and 62.2%, respectively. VBNC *V. parahaemolyticus* ATCC 33844 and *V. parahaemolyticus* ATCC 27969 exhibited the highest hydrophobicity in levels of 41.1-43.7 % in ASW_30_.

**Table 5.** Hydrophobicity (%) * of *Vibrio parahaemolyticus* int the viable but nonculturable state induced in artificial sea water (ASW, pH 6) microcosms for 100 days at 4°C.

| *V. parahaemolyticus*<br>strain | Microcosm † | Incubation period (days) |  |
| --- | --- | --- | --- |
|  |  | 0 | 100 |
| ATCC 17802 | ASW | 37.1 | ND ‡ |
|  | ASW<sub>5</sub> | 26.3 | 37.5 |
|  | ASW<sub>10</sub> | 45.8 | 46.8 |
|  | ASW<sub>30</sub> | 27.2 | 62.2 |
| ATCC 33844 | ASW | 22.0 | ND |
|  | ASW<sub>5</sub> | 22.8 | 20.9 |
|  | ASW<sub>10</sub> | 22.2 | 37.1 |
|  | ASW<sub>30</sub> | 29.8 | 41.1 |
| ATCC 27969 | ASW | 22.0 | 25.2 |
|  | ASW<sub>5</sub> | 23.0 | 17.5 |
|  | ASW<sub>10</sub> | 23.1 | 23.1 |
|  | ASW<sub>30</sub> | 20.8 | 43.7 |

\* Hydrophobicity (%) of *V. parahaemolyticus* strains was calculated, following an equation as; $(A_0 - A_1)/A_0 \times 100$ . $A_0$ , the absorbance of the untreated cell suspension at 600 nm; $A_1$ , the absorbance of the bacterial assay tube at 600 nm.
† Stationary growth phase cells of *V. parahaemolyticus* were incubated in the modified microcosms with different NaCl contents; 0.75% (ASW), 5% (ASW<sub>5</sub>), 10% (ASW<sub>10</sub>), and 30% NaCl (ASW<sub>30</sub>) at 4°C.
‡ Not determined.

### Morphological change

The pure cultures of *V. parahaemolyticus* ATCC 17802 were filled with lots of granules in cytoplasm and their cell membranes were shown to become intact without minor damages (Fig 3A). By contrast, VBNC *V. parahaemolyticus* ATCC 17802 cells had the less organized cytoplasmic layers. Particularly, cell membrane of VBNC *V. parahaemolyticus* was largely loosened, with the generation of empty gaps between the inner and the outer membranes (Fig 3B-C). Importantly, *V. parahaemolyticus* cells acquired the aberrantly-shaped coccal morphologies after the entry into the VBNC state.

**Fig 3.** Transmission electron microscopic (TEM) images of *Vibrio parahaemolyticus*. *V. parahaemolyticus* ATCC 33844 grown in tryptic soy broth supplemented with 3% NaCl at 37°C for 24 h (A) or incubated in artificial sea water (ASW; pH 6) (B) and ASW (pH 6) supplemented with 5% NaCl (C) at 4°C for 100 days.

## Discussion

After induction of VBNC forms, cell membrane integrity can be measured via epifluorescence microscopy with dual-staining of membrane permeabilizing probes such as SYTO9 and PI (20, 21). In this study, 100 days of starvation at 4°C resulted in the inability of *V. parahaemolyticus* to grow, whereas the cell number with intact membranes was consistently stable over several months of cold-starvation (Fig 1). *V. parahaemolyticus* was shown to maintain its membrane structure and integrity ranging from 4.8 to 6.5 log CFU/slide during cold-starvation, regardless of the different NaCl contents in ASW microcosms. In particular, *V. parahaemolyticus* was induced into the VBNC state in ASW microcosms supplemented with higher NaCl contents at 4°C within 21 days and persisted for 150 days in the adverse environments. Although the addition of NaCl in low acidified foods is generally known to preserve and inhibit the growth of spoilage and pathogenic microorganisms, this study may indicate that NaCl can be an important determinant that induces or accelerates the generation of VBNC cells. However, there may be a matter of debate whether *V. parahaemolyticus* cells would be still alive in microcosms supplemented with >5% NaCl for more than 100 days. *V. parahaemolyticus* is a moderate halophilic, with optimal growth at 3% NaCl (22), and some strains can grow at 9.6% NaCl (23). Alam et al. (24) determined effects of prolonged cold-starvation and biofilm formation on the viability of VBNC *Vibrio cholerae* O1. During cold-starvation, *V. cholerae* persisted in the VBNC state for 495 days. VBNC *V. cholerae* cells within biofilms were recoverable through animal passage challenge even after having been starved at 4°C for more than one year. In this way, resuscitative effects can be one possible explanation for estimating the viability of VBNC cells. After 150 days of persistence in in ASW microcosms (pH 6) at 4°C, VBNC *V. parahaemolyticus* cells were reverted to a culturable state following temperature upshift in a formulated resuscitation buffer (data not shown). Accordingly, it was found that *V. parahaemolyticus* was able to enter the VBNC state in low acidified and nutrient-deficient environments containing high NaCl concentrations at 4°C, and VBNC cells retained their membrane structure and integrity under cold-starvation conditions consistently. Until now, many studies were undertaken to investigate phase transition of microorganisms into the VBNC state caused by various environmental stresses, there is still insufficient information to determine whether the complex factors (low temperature, starvation, NaCl, and low pH) trigger the formation of VBNC cells. Thus, further studies are necessary to ensure the accurate and effective identification of VBNC bacteria, as well as their pathogenic potentials.

ROS compounds play an important role on the loss of culturability and formation of VBNC cells (1, 2, 25, 26, 27). As aerobic organisms respond to oxidative stress, major cellular components such as polyunsaturated fatty acids and proteins on membrane are directly degraded by ROS compounds. Bacteria may begin to be injured and altered at the essential site of cell membranes when the concentration of active ROS substances increases to a level that exceeds the cell’s defense capacity, thereby causing a decrease in membrane fluidity (28, 29). In this study, VBNC *V. parahaemolyticus* ATCC 17802 induced in ASW microcosms (pH 6) at 4°C for 100 days exhibited the increased catalase activities more than those for the actively growing cells (Table 3). In the VBNC forms of *V. parahaemolyticus* ATCC 33844 and *V. parahaemolyticus* ATCC 27969, the catalase activities fluctuated largely between 5.868-111.502 U mg^−1^ proteins. Although the enzymatic activities of catalase remained inconsistent, depending on the *V. parahaemolyticus* strains and NaCl concentrations, we assumed that under cold-starvation conditions, a mechanical degradation of cell membrane caused by ROS would be closely associated with the formation of VBNC *V. parahaemolyticus*. Noor et al. (26) observed that expression of *KatE* was known to be involved in encoding some protective proteins such as catalase HP II with specific functions in oxidative stress defense, and the *KatE* mutation led to significant decreases in culturable numbers of *E. coli* K-12. *E. coli* K-12 cells might be degraded by ROS due to the deletion of *KatE* and give rise to the introduction of VBNC forms.

Peculiarly, correlations between cold-starvation and membrane potential of VBNC cells were reported by several studies (13, 30). In this study, the increase in fluorescence due to partitioning of NPN uptake into outer membrane was measured by the prolonged incubation of *V. parahaemolyticus* under cold-starvation conditions. VBNC *V. parahaemolyticus* ATCC 17802 induced in ASW and ASW_5_ at 4°C for 30 days showed the increasing levels of NPN uptake more than those for the pure cultures (Table 1). When *Micrococcus luteus* was incubated in lactate (0.01%) minimal medium at 4°C, this organism became VBNC after 30 days and exerted a reduction of membrane potential, as evidenced by quantitative flowcytometry with Rhodamine 123 probe that is indicative of viable or non-viable cells (30). As well-organized in a study of Trevors et al. (14), if bacteria underwent a phase transition to the VBNC state temporarily, membrane became less fluid with an intracellular leakage (K^+^) from cytoplasm, supporting our findings. Using radioactive probe (tetra[^3^H]phenylphosphonium bromide), VBNC *C. jejuni* strains persisted in natural water at 4°C for 30 days showed dramatically reduced membrane potentials at levels of 2-14 mV (those for the stationary phase cells ranged from 54 to 79 mV) (Tholozan et al. (13). The authors also reported that 30 days of cold-starvation caused at least 10^2^-fold decreases in internal K+ concentrations of VBNC *C. jejuni* cells. Initially, the leakages of protein were 0.433-1.562 at OD_240 nm_ in ASW solutions (Table 2). After 150 days of incubation at 4°C, the high concentrations of protein leaked from VBNC. *V. parahaemolyticus* cells, ranging from 0.868 to 3.755 at OD_240 nm_. The alteration of membrane permeability may correspond to a decrease in cell membrane fluidity, as evidenced by an imbalance within the bacterial cells, which had the low membrane potentials due to the penetration of NPN probes and the leakages of cellular contents such as protein and DNA. Furthermore, when *V. parahaemolyticus* persisted for 90 days in the VBNC state at 4°C, the total amount of saturated fatty acids weas slightly increased, showing by 42.4%-47.2% (Table 4). Wong et al. (2) showed that lauric acid, myristic acid, pentadecanoic acid (_C15_), and palmitic acid were found to be increased in VBNC *V. parahaemolyticus* ST550 cells induced in MMS at 4°C for 35 days. Food-isolated strains of *V. parahaemolyticus* commonly exhibited increased concentrations of decanoic acid (_C10_), tridecanoic acid (_C13_), and myristic acid after induction of the VBNC state (10). Gram-negative bacteria typically alter their membrane fluidity with significant changes in the ratio of saturated fatty acid to unsaturated fatty acid, the levels of cyclopropane fatty acid, and *cis/trans* isomerization in response to external environmental conditions (19). As determined by Chiang et al. (31), who showed that acid-adaptation at pH 5.5 for 90 min increased the ratio of saturated fatty acid/unsaturated fatty acid in *V. parahaemolyticus* cells, the acidified microcosms used in this study would be linked to the increased concentration of saturated fatty acids in VBNC *V. parahaemolyticus* cells. An increase in the amount of palmitic acid and stearic acid was shown to be involved in increasing membrane rigidity in *V. parahaemolyticus* cells (32). Meanwhile, VBNC cells had comparatively increased hydrophobicity (14, 33). The increasing membrane rigidity would be closely involved in the maintenance of membrane integrity, thereby making the extraction of DNAs from VBNC cells more difficult.

## Conclusion

At the onset of cold-starvation, *V. parahaemolyticus* used in this study was induced into the VBNC state, while retaining its membrane integrity. The higher the NaCl concentrations, the faster is the shift into the VBNC state. *V. parahaemolyticus* underwent selected physiological changes, such as modulation of membrane potential, and re-arrangement of fatty acid composition and hydrophobicity that may result in a decrease of cell fluidity, as the cells were induced into the VBNC state during cold-starvation. Theoretically, the physiological modulations may lead to the dwarfing of *V. parahaemolyticus* cells with the flappy outer membrane out of cytoplasm, thereby minimizing their cell maintenance requirements. *V. parahaemolyticus* responds to a certain environmental stress such as cold-starvation by inducing its phase transition into a VBNC state.

## Acknowledgement

This research was supported by Basic Science Research Program through the National Research Foundation of Korea (NRF) funded by the Ministry of Science and ICT (NRF-2016R1A2B4014591 and NRF-2018R1A6A1A03025159).

## Author contributions

Jae-Hyun Yoon performed experiments, analyzed data, and write the manuscript. Jeong-Eun Hyun helped to perform experiment and revise the manuscript Sung-Kwon Moon helped to analyze data and revise the manuscript. Sun-Young Lee designed and supervised the research and co-wrote the manuscript.

